# A novel approach to derive human midbrain-specific organoids from neuroepithelial stem cells

**DOI:** 10.1101/061077

**Authors:** Anna S. Monzel, Lisa M. Smits, Kathrin Hemmer, Siham Hachi, Edinson Lucumi Moreno, Thea van Wuellen, Ronan M.T. Fleming, Silvia Bolognin, Jebs C. Schwamborn

## Abstract

Research on human brain development and neurological diseases is limited by the lack of advanced experimental *in vitro* models that truly recapitulate the complexity of the human brain. Furthermore, animal models of human neurodegenerative diseases have failed dramatically, and the success rate of clinical trials based on these models has been disappointing. Here, we describe a novel and robust human brain organoid system that is highly specific to the midbrain derived from regionally patterned neuroepithelial stem cells. These human midbrain organoids contain spatially organized groups of dopaminergic neurons, which make them an attractive model to study Parkinson’s disease. Midbrain organoids are characterized in detail for neuronal, astroglial and oligodendrocyte differentiation. Furthermore, we show the presence of synaptic connections and electrophysiological activity. The complexity of this model is further highlighted by the myelination of neurites. The present midbrain organoid system has the potential to be used for advanced *in vitro* disease modeling and therapy development.

## Introduction

With the development of methods to generate induced pluripotent stem cells (iPSCs) from somatic cells (1,2), human cells with the potential to generate all body cell types in vitro became available. This advance led to tremendous progress in the development of protocols for the differentiation of iPSCs into various human cell types. Additionally, disease-specific human iPSCs and their derived cell types are now widely used for *in vitro* disease modeling. However, particularly with regard to neuronal diseases, it is of importance to consider that the human brain is an extremely complex, three-dimensional structure. Consequently, the investigation of its development and modeling of disease processes in traditional, twodimensional cultures has strong limitations. It has been demonstrated that the presence of a 3D matrix promotes many biologically relevant functions, such as differentiation capability (3–6), cellular signaling, and lineage specification (4, 7, 8), which are otherwise not observed in 2D monolayer cell cultures (9). Additionally, impaired cell-cell and cell-matrix interactions have been shown (10). These observations have, in recent years, prompted the use of human iPSCs for the generation of three-dimensional *in vitro* models of complete organs, the so-called organoids.

These organoid technologies have been pioneered for the small intestine (11) and later extended also to other organs or parts of organs (12). Recently, protocols for the generation of human brain-like organoids have been developed, including protocols for cerebral (13), cerebellar (14) and forebrain organoids (15). These organoids provide a proof-of-concept that human iPSCs can indeed differentiate into various cell types and even self-organize with a specific spatial orientation, which recapitulates key features of the human brain. These brain organoids have been used successfully to model a genetic form of microcephaly (13) as well as Zika virus-induced microcephaly (15). Importantly, thus far, all brain organoids have been generated directly from iPSCs; however, evidence suggesting that these organoids could also be derived from more fate-restricted neural stem cells is lacking. The utilization of neural stem cells as the starting population has the advantage that already patterned cells differentiate faster and can be cultured more efficiently (cheaper, faster cell doublings, ease of handling, etc.). Furthermore, while the generation of certain brain structures, such as the cerebral cortex or cerebellum, is meanwhile well described, a higher degree of pre-patterning seems to be required for the generation of other highly specialized structures. This is particularly true for brain regions severely affected in neurodegenerative disorders, such as Alzheimer’s disease (hippocampus) or Parkinson’s disease (substantia nigra).

To address these challenges, we used our previously described human neuroepithelial stem cell (NESC) culture system (16) and differentiated these NESCs under dynamic conditions into human midbrain specific organoids.

## Results

### Generation of human midbrain-specific organoids

Previously described neuroepithelial stem cells (16) were used as the starting population for the generation of human midbrain organoids. Compared to iPSCs as a starting population, NESCs are already patterned towards midbrain/hindbrain identity. Therefore, a faster and more complete differentiation into midbrain organoids was expected.

Typically, NESCs express the neural progenitor markers SOX1, SOX2, PAX6, and NESTIN prior to organoid generation (Fig. S1A). Cells were seeded on round-bottom ultralow adhesion 96-well plates enabling the cells to form three-dimensional colonies. They were cultured in the presence of the GSK3b inhibitor CHIR99021 to stimulate the canonical WNT signaling pathway, and the SHH pathway was activated using purmorphamine (PMA). On day 8, the three-dimensional NESC colonies were embedded into droplets of Matrigel for structural support, and two days after, specification into midbrain organoids was initiated by changing the media to differentiation media. We kept the organoids in 10cm Petri dishes for short-term cultures or in ultralow adhesion 24-well plates for long-term cultures and placed them on an orbital shaker rotating at approximately 80 rpm (Fig. 1A).

**Fig. 1.**
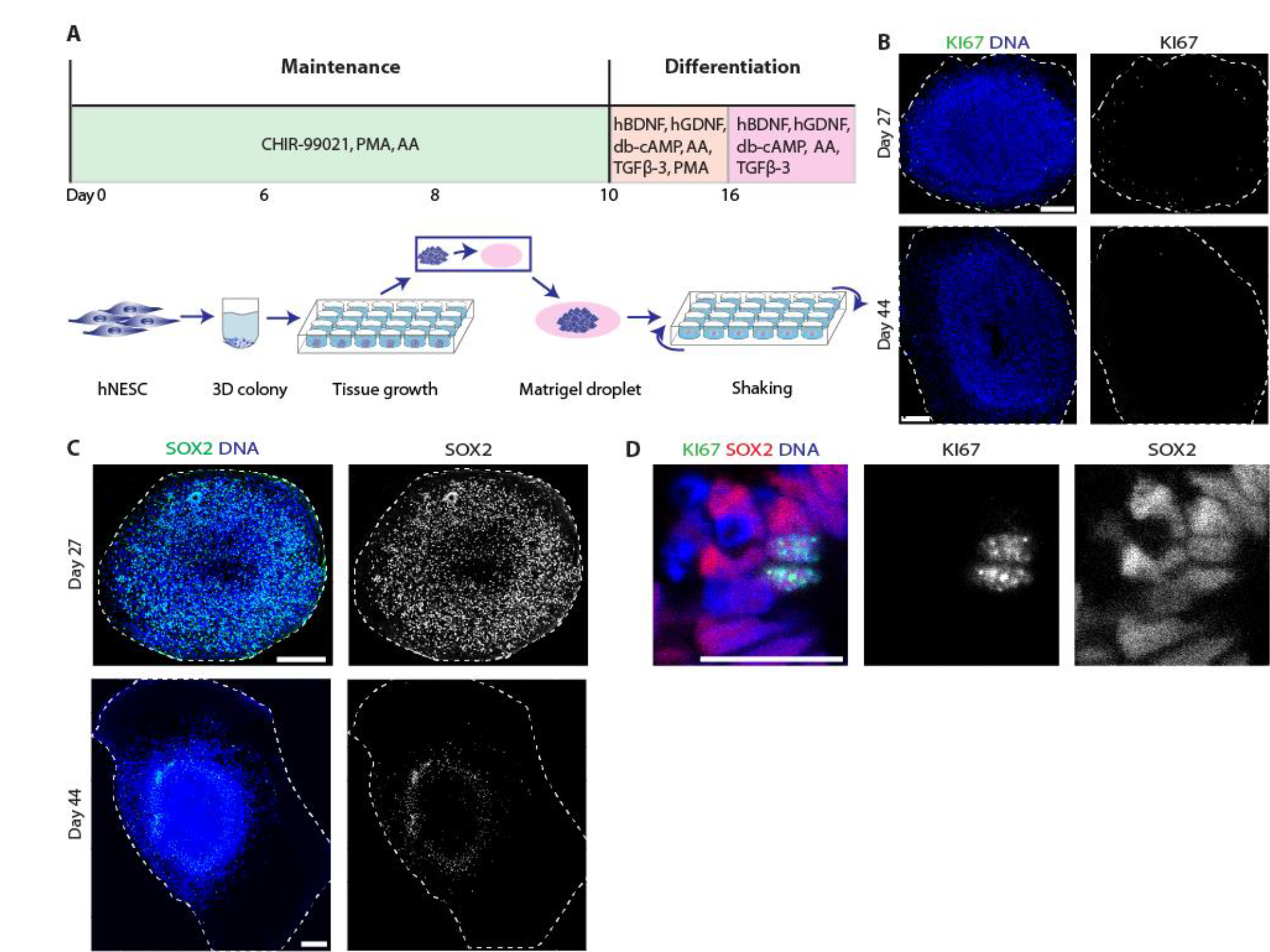
Derivation of midbrain-specific organoids from human neuroepithelial stem cells. (A) Procedure of midbrain organoid culture system. Details are described in the methods section.hNESC, human neuroepithelial stem cell; AA, ascorbic acid, PMA, purmorphamine. (B) Immunohistological staining for the cell proliferation marker Ki67 at day 27 and day 44 of the organoid culture reveals a decrease in the amount of proliferative cells in the midbrain organoids. (C) Immunohistological staining at day 27 and day 44 for the neural stem cell marker SOX2. SOX2 expression becomes more regionally restricted at the later stages. 150 μm section (C, lower panel). Scale bars, 200 μm (B, C); 20 μm (D). Dashed lines indicate the perimeter of the organoid.

At the early stages (27 days), midbrain organoids widely expressed the cell proliferation marker Ki67, which decreases upon maturation (Fig. 1B). Furthermore, young organoids expressed the neural progenitor marker SOX2, which also decreases during maturation and becomes more regionally restricted, resembling the formation of a stem cell niche (Fig. 1C). Although consistent with previous findings (13,17, we found substantial cell death in the core of the organoids, which resulted from the lack of nutrient support in the center of the organoid. In general, we did not observe high levels of apoptotic cell death, revealed by staining of the marker cleaved caspase-3 (CC3). The basic level of apoptosis that was detectable did not change during the course of differentiation (Fig. S1B). Importantly, we were able to reproduce the generation of midbrain organoids from NESCs without significant variation.

### Neuronal differentiation and self-organization of human neuroepithelial stem cell derived midbrain organoids

After showing a decrease of proliferation and stem cell identity, we assessed neuronal differentiation. Because we are particularly interested in the future utilization of midbrain organoids for in vitro modeling of Parkinson’s disease, we also investigated the specification of midbrain dopaminergic neurons (mDNs). We were able to see robust differentiation into TUJ1 positive neurons and TH positive DNs. These stainings revealed the formation of a complex neuronal network (Fig. 2A, 2B). Double-positive staining of the mature neuronal marker MAP2/TH demonstrated the maturation of DNs within the organoids (Fig. S2A). We further examined whether NESC-derived organoids undergo differentiation into DNs with midbrain identity. In late-stage organoids, a large population of TH-, LMX1A-, and FOXA2- positive neurons was observed (Fig. 2C-E). qRT-PCR further revealed the upregulation of mDN differentiation markers, including LMX1A, LMX1B, EN1, NURR1, AADC, and TH (Fig. 2F, 2G). These data indicate that the obtained dopaminergic neurons indeed have midbrain identity.

**Fig. 2.**
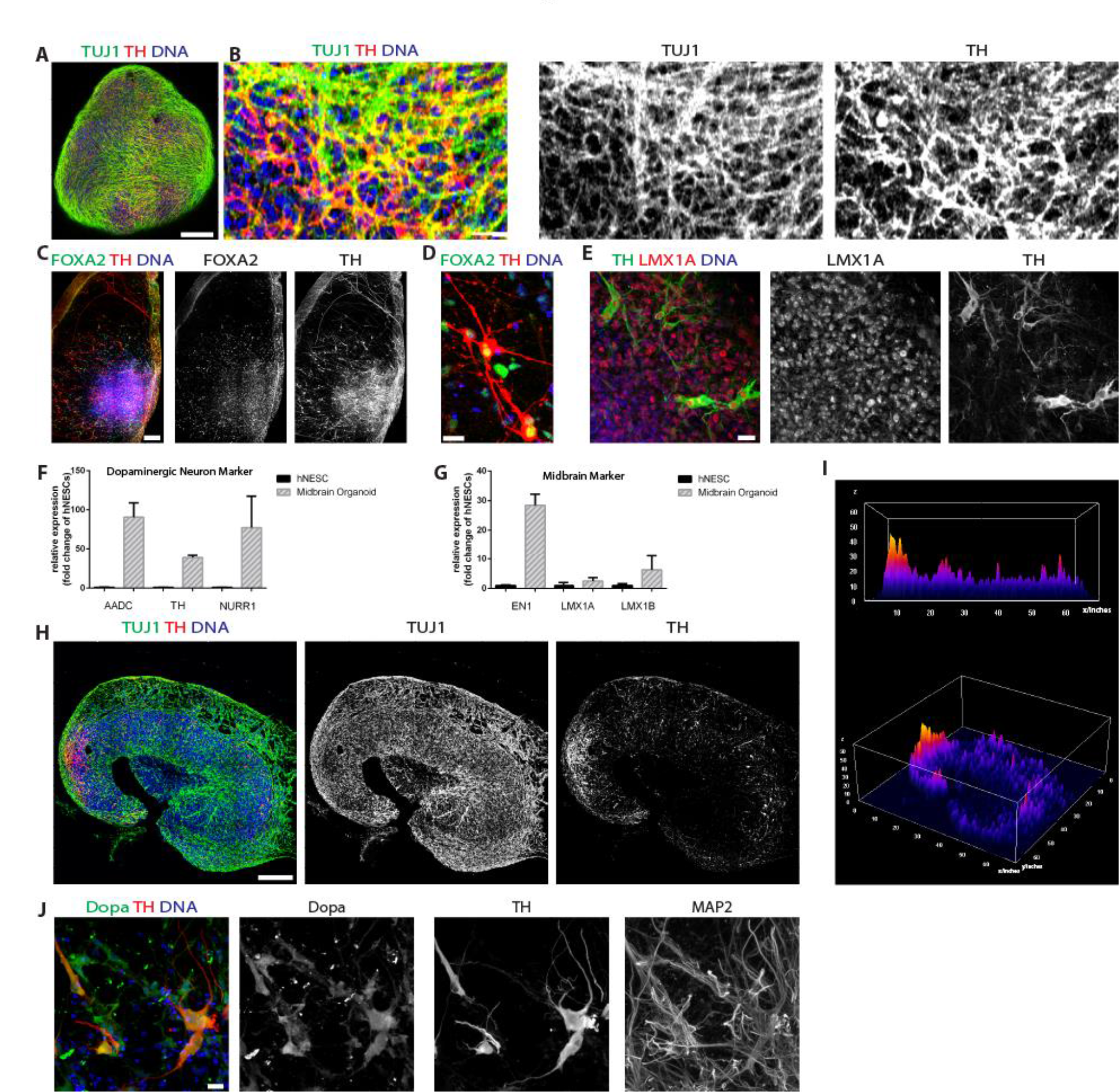
Neuronal differentiation and self-organization in midbrain organoids. (A) Whole-mount immunohistological staining of an organoid at day 27 for the DN markers TUJ1 and TH. (B) Higher magnification of (A). (C) Immunohistological staining for the mDA neuron markers FOXA2 and TH at day 61 reveals the midbrain identity of hNESC-derived organoids. (D) Higher magnification of (C). (E) High magnification of LMX1A/TH-positive mDA neurons in midbrain-specific organoids. (F-G) qRT-PCR analysis for the mDN markers AADC, TH, NURR1 (F) and EN1, LMX1A, LMX1B (G) at day 48. Error bars indicate the standard deviation from 3 independent cultures. (H-I) Asymmetry analysis of DNs. Immunostaining of DA neurons for TUJ1 and TH (H) was analyzed based on fluorescence intensities using a 3D surface plot (I). (J) Immunohistological staining for dopamine, MAP2 and TH reveals the presence of dopamine producing neurons. 150 μm sections (C, H). Scale bars, 200 μm (A, C, H), 20 μm (B, D, E, J).

In previous studies, it has been repeatedly shown that stem cells exhibit an enormous potential to self-organize into complex heterogeneous brain organoids (13, 14, 18, 19). To examine the degree of spatial organization in NESC-derived midbrain organoids, we evaluated the distribution pattern DA neuronal markers TUJ 1/TH and depicted the results using surface plots. Strikingly, we found that DA neurons form clearly specified clusters within midbrain organoids (Fig. 2H, 2I). To further demonstrate the identity of TH positive neurons as dopaminergic, we analyzed their ability to produce the neurotransmitter dopamine. Immunostainings of mature organoids demonstrated the presence of dopamine and TH-double positive cells (Fig. 2J). These results indicated that mDNs of NESC-derived organoids self-organize into a complex, spatially patterned and functional neuronal tissue.

### Glial differentiation in midbrain organoids

During the development of the fetal human brain, neural tube-derived cells not only differentiate into neurons but also into glia cells, including astrocytes and oligodendrocytes. Therefore, we investigated the presence of these glia cells in the midbrain organoids. In good agreement with brain development, where glia differentiation temporally follows neuronal differentiation, we were unable to detect significant amounts of glia cells in young organoids (day 27). However, in more mature organoids (day 61), we observed astrocytes positive for the markers S100P and GFAP. Interestingly, we obtained populations of astrocytes in both a quiescent state (negative for GFAP) and a reactive state characterized by GFAP expression (Fig. 3A, B). Moreover, we detected a fraction of cells that differentiated into O4-positive oligodendrocytes at day 44. Furthermore, interestingly, these oligodendrocytes typically showed a spatially asymmetric distribution within the organoids (Fig. S3A). In the central nervous system, mature oligodendrocytes form myelin sheaths that enwrap axons to accelerate the transmission of action potentials along axons. To analyze if the oligodendrocytes within the midbrain organoids are able to execute their actual function, i.e., formation of myelin sheets, we performed immunofluorescence staining against 2’,3’-cyclic-nucleotide 3’-phosphodiesterase (CNPase), a myelin-associated enzyme together with the neuronal marker TUJ1. A 3D surface reconstruction of these stainings revealed numerous TUJ1-positive neurites that were ensheathed by myelin sheets of CNPase-positive oligodendrocytes (Fig. 3C and S3B). Interestingly, these neurites often showed gaps of ensheathment, resembling the formation of nodes of Ranvier (Fig. 3C) that allow for saltatory fast neuronal transmission.

**Fig. 3.**
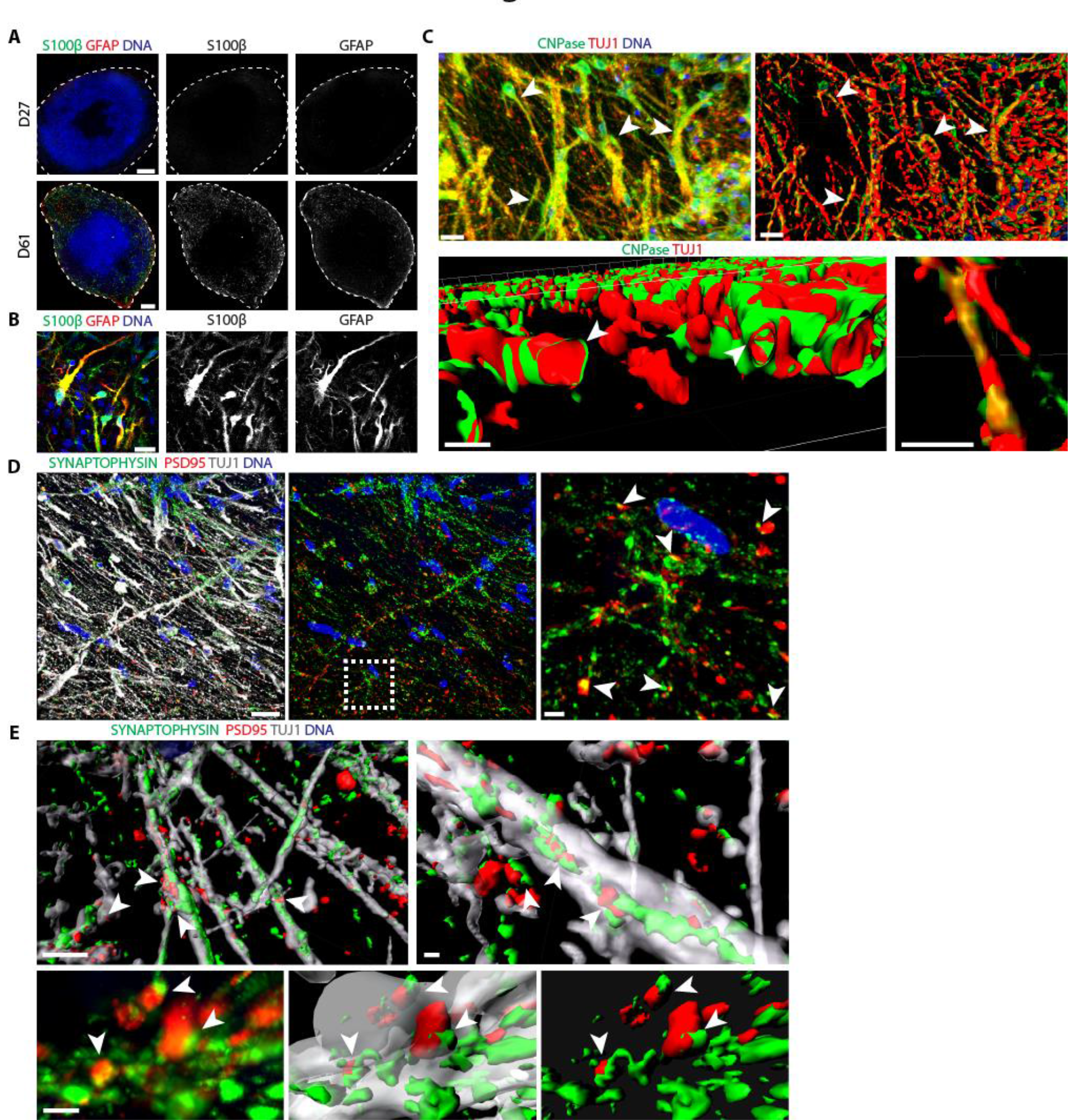
Differentiation into glia cells and formation of synaptic connections. (A) Immunohistological staining for the astrocytic markers S100P and GFAP in early-(day 27) and late-stage (day 61) organoids. Dashed lines indicate the perimeter of the organoid. (B) Higher magnification of (A) at day 61 showing astrocytes expressing S100P and GFAP. (C) Immunohistological staining of an organoid at day 61 revealing robust differentiation into CNPase-positive oligodendrocytes. Three-dimensional surface reconstructions of confocal z-stacks visualized the formation of myelin sheaths that enwrap TUJ1-positive neurites (arrowheads) as well as the formation of nodes of Ranvier that are suggested by the presence of gaps of CNPase-positive ensheathment. (E) Immunohistological staining of the presynaptic marker synaptophysin and the postsynaptic marker PSD95 at day 61. Arrowheads indicate a direct contact between a pre-and a postsynapse. Dashed box indicates the region of magnification. Images show the 3D view of a confocal z-stack. (F) Three-dimensional surface reconstructions of confocal z-stacks demonstrate the formation of synaptic connections between different neurites of an organoid as indicated by several direct contacts (arrowheads) between the pre-and postsynaptic markers synaptophysin and PSD95, respectively. Lower panels show high magnifications of a 3D view of a confocal z-stack and the corresponding 3D surface reconstruction of several synaptic connections. Scale bars, 200 μm (A), 20 μm (B, C upper panel, C lower panel left, D upper/middle panel left/middle), 2 μm (C lower panel right, D upper/middle panel right, D lower panel).

### Functionality of midbrain organoids

One important requirement for neuronal transmission is the development of a mature neuronal network via the formation of synaptic connections. Therefore, synaptic connectivity was investigated using immunohistological staining against the presynaptic marker synaptophysin and the postsynaptic marker PSD95 at day 61. A subsequent 3D surface reconstruction demonstrated not only the formation of numerous pre-and postsynaptic puncta but also multiple synaptic connections (Fig. 3D and S3C). Synaptic connections have been developed between different neurites, indicated by the direct contact of synaptophysin-positive presynapses with PSD95-positive postsynapses (Fig. 3E). Accordingly, midbrain organoids exhibit the ability to forward signals via synaptic connections and thus fulfill the prerequisite for being electrophysiologically functional.

To further confirm their functionality and neuronal network activity, we performed Fluo-4AM calcium imaging on whole organoids. We measured the spontaneous neuronal activity based on calcium transients evoked by action potentials (Fig. 4A, B, Fig S4, video S1). Notably, some of the fluorescent traces showed regular firing patterns, which were indicative of tonic electrophysiological activity and resembled the pacemaker activity of dopaminergic neurons (20,21. In addition to calcium imaging, a multielectrode array (MEA) system was used to examine the electrophysiological activity. This methodology allows non-invasive recordings of extracellular field potentials generated by action potentials. At day 60-70, the midbrain organoids were placed on a grid of 16 electrodes in a 48-well tissue culture plate (Fig. 4C). Spontaneous activity was detected over several days by individual electrodes in the form of mono- and biphasic spikes (26.12 ± 5.1 spikes/active electrode (n≥3), Fig. 4D,E). Furthermore, spikes occurred close in time on multiple electrodes, which represents neuronal network synchronicity (Fig. 4F). These findings indicate that midbrain organoids develop functional synaptic connections and show spontaneous neuronal activity.

**Fig. 4.**
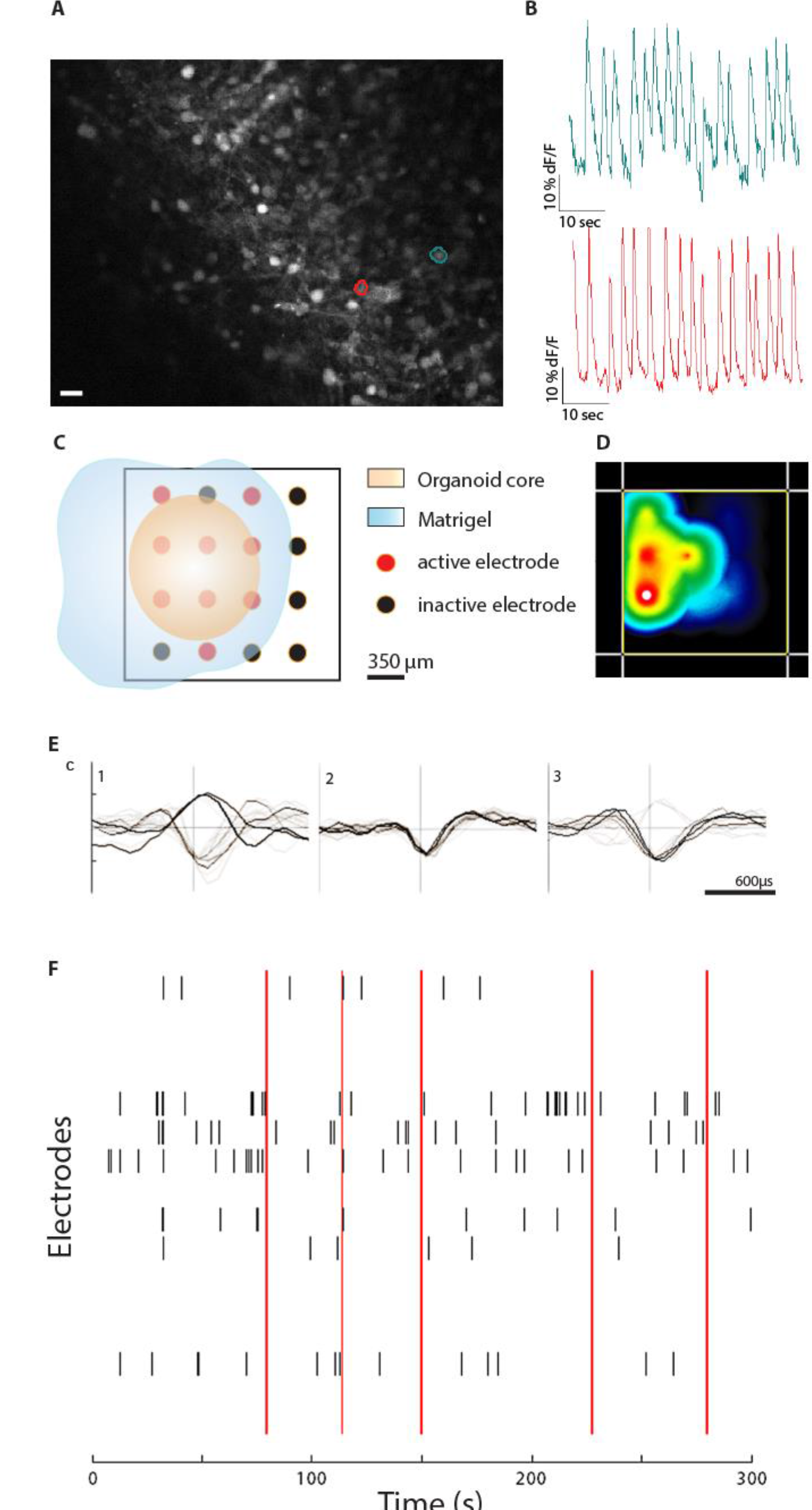
Midbrain organoids reveal electrophysiological activity. A)-B) Monitoring of the spontaneous electrophysiological activity in an organoid using Fluo-4AM based calcium imaging. A) Mean fluorescence frame of a calcium imaging dataset of a midbrain organoid with two segmented neurons expressing spontaneous activity. Scale bar, 20 μm. B) Fluorescence traces corresponding to the segmented cell bodies in (A) showing firing patterns with pacemaking-like shape. C)-F)Evaluation of the spontaneous activity in midbrain organoids after 60-70 days using a multielectrode array (MEA) system. C) Representative scheme of positioned midbrain organoid on a 16-electrode array in a 48-well tissue culture plate. D) Representative image of the activity map. E) Examples of mono-and biphasic spikes detected by individual electrodes. F) Representative image of a spike raster plot showing neuronal network activity in time and space. Spikes occurring on multiple electrodes, closely in time, represent network synchrony, indicated by red lines.

## Discussion

One of the main limitations in neuroscience and the modeling of neurological diseases is the lack of advanced experimental *in vitro* models that truly recapitulate the complexity of the human brain. Currently, although pre-clinical tests in animals suggest that a treatment will be safe and effective for a certain disease in humans, a high percentage of potential drug candidates fail in clinical trials. As an example, the failure rate for Alzheimer’s disease drug development is an astonishing 99.6% (22). Here, we have presented a novel human brain organoid system that is highly specific to the midbrain. These midbrain organoids contain spatially patterned groups of dopaminergic neurons, which make them an attractive model for the study of Parkinson’s disease. Midbrain organoids are characterized in detail with regard to neuronal differentiation and activity as well as for astroglial and oligodendrocyte differentiation.

The presence of astrocytes is crucial for the formation of synapses and regular neuronal activity (23). Astrocytes are specified later in development than neurons (24,25. Accordingly, midbrain organoids show a robust astrocyte immunoreactivity only after 61 days of differentiation. Furthermore, synaptic connections, consisting of a direct contact between pre- and post-synapses, are detectable in the midbrain organoids. These synaptic contacts are the prerequisite for electrophysiological and neuronal network functionality, which we indeed detected in the midbrain organoids by Ca2+ imaging and multi-electrode array measurements. Additionally, fast information transmission between neurons depends on axonal myelination, which is achieved by oligodendrocytes. In most stem cell based differentiation protocols, the differentiation into oligodendrocytes is extremely inefficient (26,27. However, in the present approach, we achieved a robust differentiation into oligodendrocytes and a high degree of neurite myelination. Neurites in these midbrain organoids are ensheathed by oligodendrocytes and even structures, such as the nodes of Ranvier that are of critical importance for saltatory transmission of signals in axons (28), become apparent.

Compared to other pioneering human brain organoid systems, we are able to generate organoids of a remarkable size (up to 2 mm in diameter) with a high reproducibility. Furthermore, we provide a detailed characterization of synapse formation and myelination, which has not been achieved in human brain organoids so far. Importantly, in contrast to all other human brain organoid systems, our starting population of cells are not iPSCs but neuroepithelial stem cells (16), which allows us to achieve a faster and more directed differentiation. Our approach is fully focused on the midbrain, as shown by the abundant presence of neurons with midbrain dopaminergic neuron identity. These neurons are not only able to produce dopamine but are also asymmetrically distributed in a discrete cluster. This asymmetry mirrors a unique feature of the human brain where the soma of dopaminergic neurons reside in the substantia nigra.

The presented human midbrain organoids have a great potential to be used for *in vitro* modeling of diseases that strongly affect the human midbrain, particularly Parkinson’s disease (29). The midbrain organoids may be used for investigating toxin-induced PD, e.g., after exposure to MPTP (30,31 or rotenone (31) and genetic forms of PD. In particular, the genetic forms, which are caused by mutations in genes, such as LRRK2 (32) or *Pink1* (33), could be modeled using patient-specific iPSC derived NESCs differentiated into midbrain organoids. Patient-specific midbrain organoids could reveal specific phenotypes that are not abundant in 2D cultures and therefore may be used for mechanistic studies and drug testing. Importantly, such a pipeline would be fully suitable for approaches in personalized medicine (34,35.

Neurodegenerative disorders, such as PD, are typically considered to be age-associated diseases (36,37. However, there is accumulating evidence that PD has a strong neurodevelopmental component that probably defines the susceptibility to develop the disease (38,39. This finding supports the importance of human brain development models to investigate the disease underlying mechanisms. Currently, available protocols for the generation of brain organoids typically result in structures that resemble the developing fetal human brain (13,15. Remarkably, the presented midbrain organoids exhibit a ring of neural stem cells in the center of the organoid structure, reminiscent of the stem cell niche of the ventricular/subventricular zone. Therefore, the midbrain organoids not only enable the *in vitro* modeling of PD and drug testing but also the study of the developmental aspects of the disease. To date, the development of therapies against such neurodegenerative disorders largely depends on animal models. However, over the last decades, numerous therapies that showed promising results in mice failed in clinical trials. These findings support the concept that mice are not suitable to mimic human neurodegenerative diseases. The reasons for this are numerous. As an example, a study comparing the gene expression profile of mouse models for Parkinson’s disease, Alzheimer’s disease, Huntington’s disease and amyotrophic lateral sclerosis to the corresponding human diseases revealed an extremely weak enrichment for deregulated gene sets (40). Interestingly, concerning the gene expression pattern, even healthy human aging seems to be more closely related to the named diseases than the corresponding mouse models (40). Additionally, many neurodegenerative diseases in humans simply do not occur naturally in mice. The available toxin-induced models and the genetic models are either rather artificial or have rather weak phenotypes and have not been predictive of the human response (41–43). This finding clearly implies that the underlying disease mechanisms and biology are very likely to be dramatically different between the two species.

Altogether, these findings may explain the weak performance of mouse-based translational therapies for neurodegenerative diseases and underscore that new, human specific models are strongly needed. The presented midbrain organoid system along with other models may be a first step towards a more human-specific, probably even personalized, era of advanced disease modeling and therapy development.

## Materials and Methods

### Midbrain organoid culture

The hiPSC-derived hNESCs were cultured as previously described (16). On day 0 of the organoid culture, hNESCs at passage <20 were treated with accutase for 5 min at 37°C, followed by gentle pipetting to generate single cells. A total of 9000 cells were seeded into each well of an ultra-low attachment 96-well round bottom plate (Corning) and cultured in N2B27 media supplemented with 3 μM CHIR-99021 (Axon Medchem), 0.75 μM purmorphamine (Enzo Life Science) and 150 μM ascorbic acid (Sigma) (referred to as N2B27 maintenance media). N2B27 medium consists of DMEM-F12 (Invitrogen)/Neurobasal (Invitrogen) 50:50 with 1:200 N2 supplement (Invitrogen), 1:100 B27 supplement lacking Vitamin A (Invitrogen), 1% L-glutamine and 1% penicillin/streptomycin (Invitrogen). The medium was changed every other day for 6 days, and 3D colonies were then transferred to ultra-low attachment 24-well plates (Corning) and cultured in N2B27 maintenance media.
On day 8 of the organoid culture, the 3D colonies were transferred to droplets of hESC-qualified Matrigel (BD Bioscience) as previously described (17). Droplets were cultured in N2B27 maintenance media either in 10-cm Petri dishes for short-term cultures or in ultra-low attachment 24-well plates (Corning) with one droplet per well for long-term cultures. On day 10, differentiation was initiated with N2B27 media supplemented with 10 ng/ml hBDNF (Peprotech), 10 ng/ml hGDNF (Peprotech), 500 μM dbcAMP (Peprotech), 200 μM ascorbic acid (Sigma), and 1 ng/ml TGF-β3 (Peprotech). Additionally, 1 μM purmorphamine (Enzo Life Science) was added to this medium for an additional 6 days. On day 14 of the organoid culture, the plates were placed on an orbital shaker (IKA), rotating at 80 rpm, in an incubator (5% CO2, 37°C) and the organoids were kept in culture with media changes every second or third day.

### Immunohistochemical analysis

Organoids were fixed with 4 % paraformaldehyde overnight at RT and washed 3× with PBS for 1 h. Afterwards, they were embedded in 3 % low-melting point agarose in PBS and incubated for 15 min at 37 °C, followed by 30 minutes incubation at RT. The solid agarose block was covered with PBS and kept overnight at 4 °C. If not indicated 7otherwise, 50 μm sections were cut using a vibratome (Leica VT1000s), and sections were permeabilized with 0.5 % Triton X-100 in PBS and blocked in 2.5 % normal goat or donkey serum with 2.5 % BSA, 0.1 % Triton X-100, and 0.1 % sodium azide. Sections were incubated on a shaker for 48-72 h at 4°C with primary antibodies in the blocking buffer at the following dilutions: rabbit anti-TH (1:1000, Abcam), chicken anti-TH (1:1000, Abcam), rabbit an ti-TH (1:1000, Santa Cruz Biotechnology), goat anti-SOX2 (1:200, R&D Systems), rabbit anti-SOX2 (1:100, Abcam), goat anti-SOX1 (1:100, R&D Systems), mouse anti-nestin (1:200, BD), mouse anti-Ki67 (1:200, BD), rabbit anti-CC3 (1:200, Cell Signalling), mouse anti-FOXA2 (1:250, Santa Cruz Biotechnology), rabbit anti-LMX1A (1:200, Abcam), chicken anti-GFAP (1:1000, Millipore), mouse anti-S100p (1:1000, Abcam), mouse anti-TUJ1 (1:600, Covance), rabbit anti-TUJ1 (1:600, Covance), chicken anti-TUJ1 (1:600 Millipore), rabbit anti-PAX6 (1:300, Covance), mouse anti-synaptophysin (1:50 Abcam), rabbit anti PSD-95 (1:300, Invitrogen), mouse anti-MAP2 (1:200, Millipore), mouse anti-CNPase (1:200, Abcam), and mouse anti-O4 (1:400, Sigma). After incubation with the primary antibodies, sections were washed three times in 0.05 % Triton X-100 and blocked for 30 min at RT on a shaker, followed by incubation with the secondary antibodies in 0.1% Triton X-100 (1:1000). All secondary antibodies (Invitrogen) were conjugated to Alexa Fluor fluorochromes. Dopamine was detected using a STAINperfect Immunostaining Kit (ImmuSmol) according to manufacturer’s protocol. Sections were costained with chicken anti-TH primary antibody (Abcam), and nuclei were counterstained with Hoechst 33342 (Invitrogen). Sections were mounted in Fluoromount-G mounting medium (Southern Biotech) and analyzed with a confocal laser scanning microscope (Zeiss LSM 710). Images were further processed with Zen Software (Zeiss) and ImageJ. Three-dimensional surface reconstructions of confocal z-stacks were created using Imaris software (Bitplane). The asymmetric distribution of DNs was assessed based on fluorescence intensities with the ImageJ Interactive 3D surface plot plugin.

### Quantitative real-time PCR

Total RNA was isolated from 48 days-old organoids. Typically, five organoids were pooled for one isolation. For dissociation, the organoids were washed once with PBS and lysed with QIAzol lysis reagent (Qiagen), passed through a needle three times and homogenized with QIAshredder columns (Qiagen). RNA was isolated using the RNeasy Mini Kit (Qiagen) according to the manufacturer’s instructions. Subsequently, isolated RNA was reverse-transcribed following the protocol of the High Capacity RNA to DNA Kit (Thermo Fisher Scientific). Quantitative real-time polymerase chain reactions (qRT-PCRs) were conducted with the Maxima SYBR Green qPCR Master Mix (Thermo Scientific). Amplification of 1 μg cDNA was performed in an AriaMx Real-time PCR System (Agilent Technologies) as follows: An initial denaturing step, 10 min at 95 °C, 40 cycles of denaturation for 15 s at 95 °C, annealing for 30 s at 60 °C, and elongation for 30 s at 72 °C. The expression levels were normalized relative to the expression of the housekeeping gene *RPL37A* using the comparative Ct-method 2^-ΔΔCt^. To evaluate the expression patterns of midbrain organoids, the values were compared to the expression levels of hNESCs, which were set to 1. The quality of the PCR products was assessed by melting curve analysis.

### Evaluation of electrophysiological activity

Calcium imaging and multielectrode array (MEA) recording was used to analyze the spontaneous activity of organoids at day 50-52 and day 60-70, respectively. A concentration of 5 μM cell permeant Fluo-4 AM (Life Technologies) in a neurobasal medium was added to the well and incubated for 45 min at 37 °C on an orbital shaker. Fluorescent images were acquired using a live cell spinning disk confocal microscope (Zeiss) equipped with a CMOS camera (Orca Flash 4.0, Hamamatsu). Calcium time-series were acquired at 5 Hz for approximately 2 min and stored as single images. These images were analyzed using the ADINA toolbox(44), which is publicly available software that has been developed to automatically segment individual cell bodies and separate the overlapping ones. Fluorescent traces, expressed as relative changes in the fluorescence intensity (ΔF/F), were then measured for segmented cell bodies.

MEA recording was conducted using the Maestro system from Axion BioSystems. A 48-well MEA plate containing a 16-electrode array per well was precoated with 0.1 mg/ml poly-D-lysine hydrobromide (Sigma-Aldrich) and subsequently coated with 10 μg/ml laminin (Sigma-Aldrich) for 1 h at room temperature (RT). Midbrain organoids were placed onto the array after day 60-70. A coverslip was placed on top to ensure the contact of the free floating organoid with the electrodes. Spontaneous activity was recorded at a sampling rate of 12.5 kHz for 5 min for up to five days at 37 °C in neuronal maturation media. Using Axion Integrated Studio (AxIS 2.1), a Butterworth band pass filter with 200-3000 Hz cutoff frequency and a threshold of 6 × SD were set to minimize both false-positives and missed detections. The Neural Metric Tool (Axion BioSystems) was used to analyze the spike raster plots. Electrodes with an average of ≥5 spikes/min were defined as active. The spike count files generated from the recordings were used to calculate the number of spikes/active electrode/measurement. Further details regarding the MEA system were previously described (45).

## Supplementary Materials

Fig. S1: Derivation of midbrain-specific organoids from human neurepithelial stem cells

Fig. S2: Maturation of midbrain dopaminergic neurons

Fig. S3: Robust differentiation into glia cells and formation of synaptic connections

Fig. S4: Midbrain organoids reveal electrophysiological activity

Video S1: Calcium time series of a midbrain organoid at day 52.

## Acknowledgements

The JCS lab is supported by the Fonds National de la Recherche (FNR) and by a University Luxembourg Internal Research Project grant.ASM, LMS, SH and ELM are supported by fellowships from the FNR (AFR, Aides a la Formation-Recherche). This is an EU Joint Programme – Neurodegenerative Disease Research (JPND) project.

